# DNP-assisted solid-state NMR enables detection of proteins at nanomolar concentrations in fully protonated cellular environments

**DOI:** 10.1101/2023.02.20.529239

**Authors:** Whitney N. Costello, Yiling Xiao, Frederic Mentink-Vigier, Jaka Kragelj, Kendra K. Frederick

**Author notes:** To whom correspondance should be addressed: Kendra K. Frederick **Email:**.

## Abstract

With the sensitivity enhancements conferred by dynamic nuclear polarization (DNP), magic angle spinning (MAS) solid state NMR spectroscopy experiments can attain the necessary sensitivity to detect very low concentrations of proteins. This potentially enables structural investigations of proteins at their endogenous levels in their biological contexts where their native stoichiometries with potential interactors is maintained. Yet, even with DNP, experiments are still sensitivity limited. Moreover, when an isotopically-enriched target protein is present at physiological levels, which typically range from low micromolar to nanomolar concentrations, the isotope content from the natural abundance isotopes in the cellular milieu can outnumber the isotope content of the target protein. Using isotopically enriched yeast prion protein, Sup35NM, diluted into natural abundance yeast lysates, we optimized sample composition we find that modest cryoprotectant concentrations and fully protonated environments support efficient DNP. We experimentally validated theoretical calculations of the limit of specificity for an isotopically enriched protein in natural abundance cellular milieu. We establish that, using pulse sequences that are selective for adjacent NMR-active nuclei, proteins can be specifically detected in cellular milieu at concentrations in the hundreds of nanomolar. Finally, we find that maintaining native stoichiometries of the protein of interest to the components of the cellular environment may be important for proteins that make specific interactions with cellular constituents.

## INTRODUCTION

Biological processes occur in complex environments containing a myriad of potential interactors. In-cell NMR allows us to obtain atomic resolution information of proteins in their native environments.(Selenko, Serber et al. 2006, Inomata, Ohno et al. 2009, Sakakibara, Sasaki et al. 2009, Banci, Barbieri et al. 2013, Theillet, Rose et al. 2013, Freedberg and Selenko 2014, Burmann, Gerez et al. 2020) Magic angle spinning (MAS) solid-state NMR is particularly useful to study proteins and their interactions inside cells.(Gupta, Lu et al. 2016, Kaplan, Narasimhan et al. 2016, Qiang, Yau et al. 2017, Albert, Gao et al. 2018, Narasimhan, Scherpe et al. 2019, Scherpelz, Wang et al. 2021) With the sensitivity gains conferred by dynamic nuclear polarization (DNP), MAS NMR has the sensitivity to detect proteins at very low concentrations in complex biological environments.(Frederick, Michaelis et al. 2015, Albert, Gao et al. 2018, Costello, Xiao et al. 2019, Narasimhan, Scherpe et al. 2019, Bertarello, Berruyer et al. 2022) This opens up the possibility of investigation of proteins at their native contexts and with their native stoichiometries relative to potential interactors. While the endogenous concentration of some abundant proteins can exceed a hundred micromolar(Nollen and Morimoto 2002), most proteins are present at concentrations between 10 nM and 10 µM in yeast cells and between 1 nM and 1 µM in cultured mammalian cells(Wang, Weiss et al. 2012). With the development of both the instrumentation and polarization agents for DNP continually improving the sensitivity of DNP NMR, detection of more and more proteins at their endogenous concentrations, where native stoichiometries are maintained, will potentially become possible.

The effectiveness of DNP-enhanced MAS NMR is critically dependent on sample composition and experimental conditions. Early biological DNP NMR investigations were optimized on purified samples at high concentrations (Hall, Maus et al. 1997, van der Wel, Hu et al. 2006). Because DNP NMR experiments are typically performed at low temperatures, biological samples are cryoprotected to avoid ice crystal formation and ensure homogenous dispersion of the polarization agent throughout the sample. In early work, samples of biomolecules were often cryoprotected by adding 60% *d_8_*-glycerol (v/v), because much of the early biological DNP work was on the light-sensitive protein bacteriorhodopsin (Rosay, Zeri et al. 2001, Rosay, Lansing et al. 2003, Bajaj, Hornstein et al. 2007) and 60:40 glycerol:water mixtures form optically-clear glasses at all freezing rates(Lane 1925, Inaba and Andersson 2007). Moreover, because polarization transfer from the early generation of polarization agents like TEMPO(Bajaj, Farrar et al. 2003) and TOTAPOL(Song, Hu et al. 2006) was most efficient in per-deuterated settings, the DNP matrix was optimized to have a buffer protonation level of 10% (v/v).(Rosay 2001). Even though many different matrix compositions can support glass formation(Tran, Mentink-Vigier et al. 2020) and polarization transfer mechanisms from the newer generation of polarization agents do not necessarily require sample per-deuteration (Lund, Casano et al. 2020, Harrabi, Halbritter et al. 2022), the matrix composition of 60:30:10 *d_8_*-glycerol:D_2_O:H_2_O, which is often referred to as “DNP juice”, remains widely and successfully used for DNP NMR investigations of purified biological molecules (Heiliger, Matzel et al. 2020, Zhai, Lucini Paioni et al. 2020, Elathram, Ackermann et al. 2022).

Despite broad usage, DNP juice may not support the highest possible experimental sensitivity, particularly for heterogenous samples. With the 60:30:10 composition, most of the volume of the sample is made up of glycerol, which decreases the rotor fill factor. Moreover, high per-deuteration reduces the number of protons at exchangeable sites and decreases absolute sensitivity. For example, optimization of the lipids and the type of cryoprotectant used for DNP NMR of membrane protein samples doubled DNP efficiency(Liao, Lee et al. 2016). Likewise, modern polarization agents generate faster polarization transfer, via stronger electron-electron couplings (Mentink-Vigier, Vega et al. 2017) and are thus less sensitive to the protonation level of the matrix (Harrabi, Halbritter et al. 2022). Because cellular milieu has different glassing properties than concentrated samples of purified isolated proteins, high concentrations of cryoprotectants may not be necessary. Doing so would increase the rotor fill factor and dramatically increase experimental sensitivity. Moreover, because cellular systems do not always tolerate deuteration well(Misra 1967), avoiding per-deuteration may not only increase the experimental sensitivity at exchangeable sites but also better maintain the biology integrity of the sample. Thus, modifying the composition of the DNP matrix for multi-component samples can potentially dramatically increase experimental sensitivity beyond the current state of the art and because these experiments are sensitivity limited it is important to do so.

Finally, most samples for modern biological NMR are isotopically enriched and when a sample contains a mixture of isotopically enriched and un-enriched sites, the contribution of the signal from natural abundance isotopes (at 1.1% of carbons and 0.37% of nitrogen) to the NMR spectra can typically be ignored (Schubeis, Luhrs et al. 2015, Gupta and Tycko 2018). While the sensitivity enhancements conferred by DNP are theoretically sufficient to enable detection of proteins at their endogenous concentrations (van der Zwan, Riedel et al. 2021), the increase in experimental sensitivity conferred by DNP is also sufficient to enable the study biomolecules containing only naturally abundant NMR active isotopes (Takahashi, Lee et al. 2012, Mollica, Dekhil et al. 2015, Märker, Paul et al. 2017, Kang, Kirui et al. 2019). Thus, for samples of an isotopically labeled protein present at its endogenous concentration in concentrated cellular milieu, it is possible, and even likely, that the number of NMR active isotopes present in the sample due to natural abundance will outnumber those in the isotopically enriched protein. Most solid state NMR studies of isotopically enriched proteins in native biological settings have focused on proteins present at concentrations in the high tens to hundreds of micromolar, where contributions from the natural abundance proteins in the biological setting have not been problematic (Renault, Tommassen-van Boxtel et al. 2012, Kaplan, Narasimhan et al. 2016, Narasimhan, Scherpe et al. 2019). A study of the yeast prion protein, Sup35NM, at endogenous levels in a biological setting at a concentration of 1 µM used isotopically depleted cellular milieu to eliminate the signals from natural abundance components that otherwise contributed to the spectra (Frederick, Michaelis et al. 2015). However, with the improvements in both instrumentation(Berruyer, Björgvinsdóttir et al. 2020, Harrabi, Halbritter et al. 2022) and development of more efficient polarization agents (Sauvee, Rosay et al. 2013, Harrabi, Halbritter et al. 2022), the characterization of proteins at even lower concentrations is experimentally tractable. Knowing the analyte concentrations that can theoretically be specifically detected over the natural abundance molecules in the sample is critical for interpretation.

In the study of Sup35NM, structural changes occurred in a region of the protein that is known to influence biological activity but is intrinsically disordered in purified samples. In that study, isotopically enriched Sup35NM was added to yeast lysates at endogenous levels, maintaining native stoichiometries with potential interactors. (Frederick, Michaelis et al. 2015) Maintaining the relative stoichiometries of Sup35 to the chaperones proteins is critical to retain inheritance of this functional prion from mother to daughter cell.(Allen, Wegrzyn et al. 2005, Masison, Kirkland et al. 2009, Helsen and Glover 2012, Kiktev, Patterson et al. 2012) However, it is unclear if maintaining the native stoichiometry of the protein of interest to the components of the cellular environment is an important experimental consideration. Uncertainty about this point complicates interpretation of the data of in-cell experiments, limiting the utility of that information. Here, we work with Sup35NM system because yeast tolerate growth in per-deuterated environments. Moreover, working in minimally diluted pelleted lysed cells – the final macromolecular concentration in these samples is lower than that in intact cells by about a third - enables control over sample composition. Finally, prior work in this system revealed that a region of Sup35NM that is disordered in purified settings becomes structured when the protein is investigated at its endogenous levels in biological settings.(Frederick, Michaelis et al. 2015, Costello, Xiao et al. 2019) To optimize the composition of the DNP matrix to maximize sensitivity, we altered the cryoprotectant content and amount of per-deuteration the DNP matrix and cellular milieu. To determine the limits of specificity in systems of isotopically enriched proteins in concentrated mixtures of biological molecules we validate theoretical calculations with experimental measurements. Working at a concentration of 25 µM Sup35NM that is an order of magnitude over the endogenous level, we examine the effect of altering native stoichiometries on a protein whose structure is influenced by biological environments.

## RESULTS

### Moderate concentrations of glycerol result in the best DNP performance

Because the DNP enhancements that increase experimental sensitivity depend on sample composition, we assessed DNP performance of samples of proteins diluted in yeast lysates made with different amounts of cryoprotectant as well as different deuteration levels of both the buffer and the components of the cellular lysates. To determine the optimal amount of cryoprotectant for proteins at low concentrations in cellular lysates, we prepared samples with different amounts of the cryoprotectant glycerol and measured the DNP enhancements. Because high enhancements do not necessarily correlate with the highest absolute sensitivity, we also determined the absolute signal intensity for the protein component. To do so, we collected ^15^N-filtered zTEDOR spectra which select for adjacent ^13^C-^15^N pairs which are over-represented in the added Sup35NM protein (Jaroniec, Filip et al. 2002). Samples containing 15% glycerol (v/v) had the highest DNP enhancements. The DNP enhancements for lysate samples prepared without any cryoprotection were 30 ± 10% smaller while the DNP enhancements of samples that contained 60% glycerol were 50 ± 10% smaller than the DNP enhancements of samples that contained 15% glycerol (Figure 1B, black). The absolute signal intensity for Sup35NM, as determined by ^15^N-filtered zTEDOR spectra, also occurs at 15% glycerol (Figure 1B, red).

**Figure 1.**
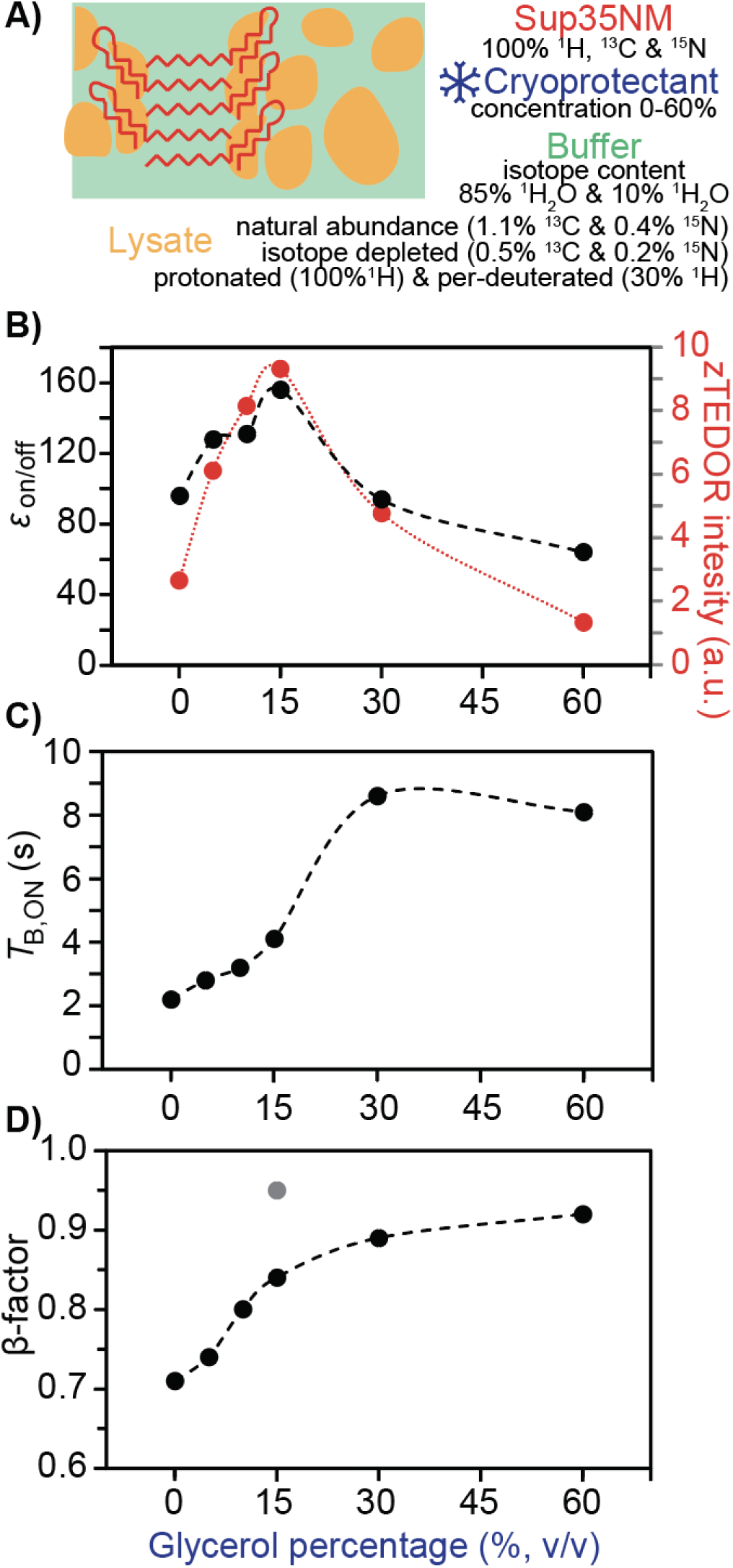
Lower cryoprotectant percentages result in better DNP performance of dilute proteins in pelleted cellular lysates. A) Sup35NM (red) is diluted in a matrix composed of cryoprotectant (blue), solvent in the buffer (green) and cellular components (yellow), all of which can have different concentrations and isotope content. (B) DNP enhancement, ε_on/off_ (black) and raw signal intensity from Sup35NM (red) using zTEDOR (^15^N-^13^C) of the carbonyl peak are both dependent upon the percentage of glycerol in the sample. Collected with a recycling delay of 4 s. (C) The build-up time values, T_B,ON_, derived from ^1^H-saturation recovery experiments fit to a stretched exponential function at the carbonyl are dependent upon the percentage of glycerol in the sample. (D) The β-factor from the ^1^H-saturation recovery experiments fit to stretched exponential decrease upon the percentage of glycerol for pelleted lysates (black) and complete lysates (gray). Samples in perdeuterated natural abundance lysates and buffer with 10% H_2_O. Dotted lines are to guide the eye. Data from one out of three independent sets of samples are shown.

We next determined build up times (*T*_B,ON_) of the yeast lysate components by fitting the carbonyl carbon (∼175 ppm) in the ^13^C-CP spectra to a stretched exponential function. If the concentration distribution of AMUPol is homogenous, the exponential will not be stretched, and β will equal 1. If the concentration distribution of AMUPol is heterogenous, there will be a mixture of underlying *T*_B,ON_ values, the increase in the range of *T*_B,ON_ values are best fit with more stretch and the value of β decreases (Rankin, Trebosc et al. 2019)(Pinon et al., 2017) (Figure 1C). Although these samples all contained similar amounts of AMUPol, the *T*_B,ON_ increased with cryoprotectant concentration from 2 seconds for samples without glycerol to 8 seconds for samples containing 60% glycerol (Figure 1C). Interestingly, the β-factor also increased from 0.75 for samples without glycerol to 0.93 for samples containing 60% glycerol (Figure 1D). Glycerol improved the dispersion of the radical throughout this complicated system; the covariance of *T*_B,ON_ with β-factor suggested that the short *T*_B,ON_ values for samples with smaller amounts of glycerol results from inhomogeneous dispersion of the radical in these sample. However low cryoprotectant percentage are not responsible for short values of *T*_B,ON_ because the *T*_B,ON_ of proline frozen with different amounts of glycerol did not show a decrease in the β-factor (β-factor = 0.97). This suggests that the change in *T*_B,ON_ values in lysate samples results from inhomogeneous dispersion of the radical and that glycerol improved the dispersion of the radical throughout this complicated system although at the expense of sensitivity and *T*_B,ON_. To determine if inhomogenous radical dispersion was a general property of cellular milieu or if it was a result of the lysate sample preparation, we prepared cellular lysates using an alternative method. We prepared lysates where the small molecule component of the sample was not removed by centrifugation before addition of 15% glycerol. We found that the fit of the build-up time for these lysates had a β-factor of 0.95, indicating that the inhomogeniety for the lysate samples resulted from the sample preparation (Figure 1D, gray point) and is not a general property of radical dispersion in cellular milieu with low concentrations of cryoprotectants. Taken together, samples containing 15% glycerol had the highest enhancements and, because of the improved fill factor, the highest overall sensitivity as well. Using 15% glycerol rather than 60% glycerol as a cryoprotectant doubles the experimental sensitivity (Figure 1B, red).

AMUPol is a highly efficient, water soluble, commercially available polarization agent. While typically used in per-deuterated environments, efficient polarization transfer from AMUPol may not require per-deuteration, even at high fields. To assess this, we first determined the dependence of DNP enhancement and *T*_B,ON_ on increasing protonation content for a standard sample of uniformly isotopically labeled proline in 60/40 glycerol/water mixtures (Figure S1). The DNP enhancement was highest when the sample contained 10% H_2_O (*v/v*) and decreased by ∼30% in a fully protonated setting while the *T*_B,ON_ values were longer by a second (*p* < 0.05, student’s t-test, n=4) (Figure S1A). To determine if analyte concentration affected DNP performance, as assessed by the magnitude of the DNP enhancement and the *T*_B,ON_ values, we varied the proline concentration by five orders of magnitude and measured DNP enhancements and *T*_B,ON_ values. Neither the DNP enhancement or *T*_B,ON_ value were sensitive to analyte concentration (Figure S1B). Thus, the DNP performance (DNP enhancement and *T*_B,ON_ values) of AMUPol is modestly improved by high deuteration levels of the buffer.

### Fully protonated environments support efficient DNP performance

Because per-deuteration is not always well tolerated by cellular systems (Misra 1967), we determined the DNP enhancement and *T*_B,ON_ values for proteins at low concentrations in cellular lysates when AMUPol is used as a polarization agent with a variety of deuteration levels. To determine if per-deuteration of the cellular lysate is required for efficient polarization transfer from the polarization agent, AMUPol, to the isotopically labeled protein of interest in samples containing cellular lysates and cryoprotected with 15% glycerol, we varied both the protonation level of the DNP sample buffer as well as the cellular lysates. First, we tested the effect of DNP sample buffer protonation levels on DNP sample performance while holding the per-deuteration level of the cellular lysates constant. We tested DNP sample buffer protonation levels of 10% and 85%. To do so, we prepared three independent DNP NMR samples of deuterated cellular lysates with 25 µM uniformly ^13^C, ^15^N-labeled Sup35NM protein, 15% *d_8_*-glycerol, 5 mM AMUPol biradical (Figure S2) and measured DNP enhancement and build up times (Figure 2, right). We found the DNP enhancement did not change (Figure 2A, right) but the build-up time for the sample containing 10% H_2_O was several seconds shorter than it was for the sample containing 85% H_2_O (Figure 2B, right). This suggested that DNP enhancements when using AMUPol as the polarizing agent in lysate samples are not sensitive to the protonation level of the DNP sample buffer though overall DNP performance is improved through the shorter *T*_B,ON_ values in protonated systems.

**Figure 2.**
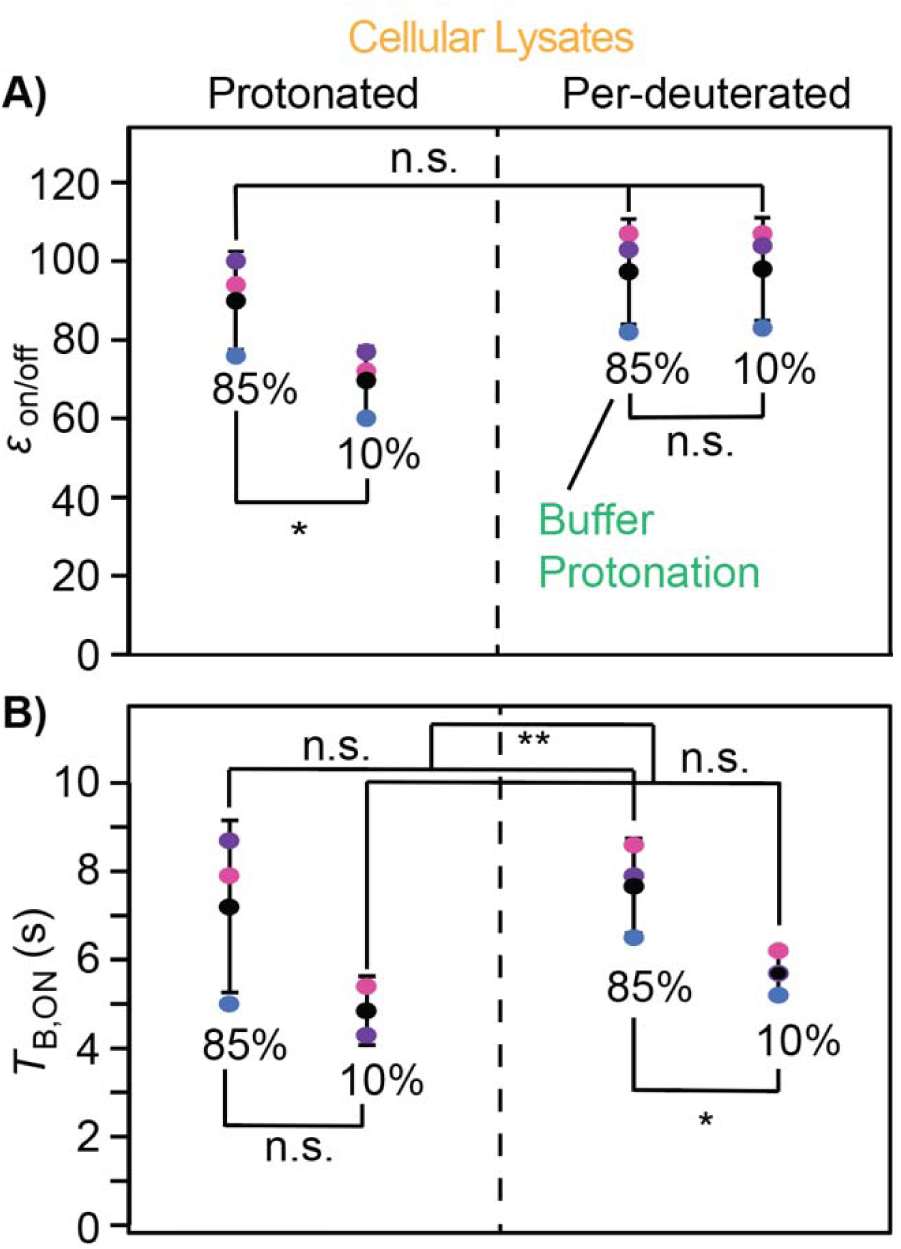
DNP performance is maintained in protonated DNP matrix conditions for cellular lysate samples using AMUPol. DNP sample buffer protonation scheme is marked with 85% or 10%. DNP cellular lysate protonation scheme is marked with protonated (100%) or per-deuterated (50%). Samples from the same data sets are indicated by color (pink, purple and blue). Average and standard deviations are shown in black. Black lines indicate results of paired student t-tests. (A) DNP enhancement (ε_on/off_) of cellular lysate samples measured by the ratio of on/off microwaves signal of the ^13^C-CP signal at the carbonyl. (B) Build-up time values derived from ^1^H-saturation recovery experiments fit to a stretched exponential function at the carbonyl are dependent upon the protonation of the buffer. Samples contain 15% d_8_-glycerol. Brackets indicate results of paired two-tailed homoscedastic student’s t-tests (n.s. p > 0.05, * p < 0.05, ** p < 0.01).

To determine if per-deuteration of the biological components of the cellular lysates improved DNP performance, we prepared DNP sample sets that contained fully protonated cellular lysates and compared the DNP enhancement and *T*_B,ON_ with the per-deuterated lysates. We found that the DNP enhancement for samples of fully protonated lysates with 10% protonated DNP sample buffer were smaller than samples with all other combinations of protonation levels of the lysate and buffer by ∼30% (Figure 2A, Student’s t-test, paired, *p* < 0.05, n=3), although the lower intensity for the sample made with 10% protonated buffer and perdeuterated lysates was a trivial result of a larger off-signal in the presence of protonated lysates that results from higher overall protonation and a shorter *T*_B,ON_. The absolute sensitivity of these experiments as assessed by the microwave on-signal intensities were similar regardless of the deuteration level of the lysate. We also found that samples with a proton content of 85%, rather than 10%, had longer *T*_B,ON_ values (Figure 2B, Student’s t-test paired, *p* < 0.001, n=6). This difference was not a result of differences in radical dispersion between protonated and deuterated sample because the β-factors were not affected by deuteration level (Student’s t-test, paired, *p* = 0.25, n=6, data not shown). However, when we probed for the uniformly isotopically enriched, fully protonated Sup35NM using ^15^N-filtered experiments, we found that the *T_B,ON_*values were not sensitive to sample deuteration levels (Student’s t-test, paired, *p* = 0.49, *n*=6, Figure S3), therefore the deuteration level modestly affects the spectroscopy of the cellular milieu but does not alter the spectroscopy of the protein of interest. Taken together, we found that highly protonated environments have modest effects on the DNP performance of AMUPol in both glycerol/water matrices and matrices containing cellular lysates. This indicates that good DNP performance from AMUPol can be expected for per-deuterated as well as fully protonated biological systems.

### Isotopically enriched proteins can be specifically detected at nanomolar concentrations in cellular milieu

The sensitivity enhancements that can be obtained for proteins diluted in biological milieu are sufficient to easily detect isotopes present at natural abundance in the sample. Thus, we calculated the expected selectivity ratio for a uniformly isotopically enriched protein diluted in yeast cellular lysates containing isotopes at their natural abundance. To do so, we determined the amount of ^13^C labeled carbonyl carbons in the 30 kDa uniformly isotopically enriched protein of interest, Sup35NM, at a concentration of 25 µM and the amount of ^13^C labeled carbonyl carbons present from natural abundance in the protein component of yeast lysates. The calculated amount of ^13^C labeled carbonyl carbon in the uniformly isotopically labeled Sup35NM at 25 µM and the calculated amount of ^13^C labeled carbonyl carbon in natural abundance yeast lysates were approximately equivalent when directly detecting ^13^C (e.g. 13C CP)(Figure 3, dashed black line). Due to the lower natural abundance of ^15^N, the calculated amount of ^15^N labeled protein backbone nitrogen in the Sup35NM was two-fold larger than that of ^15^N labeled protein backbone nitrogen in the yeast lysate(Groves, Falson et al. 1996, Frederick, Michaelis et al. 2015, Łabędź, Wańczyk et al. 2017, Oftadeh, Salvy et al. 2021). Thus, in directly detected ^13^C or ^15^N NMR experiments, the signals from natural abundance in the yeast lysate are predicted to account for half of the ^13^C carbonyl signals and a third of the ^15^N backbone signal.

**Figure 3:**
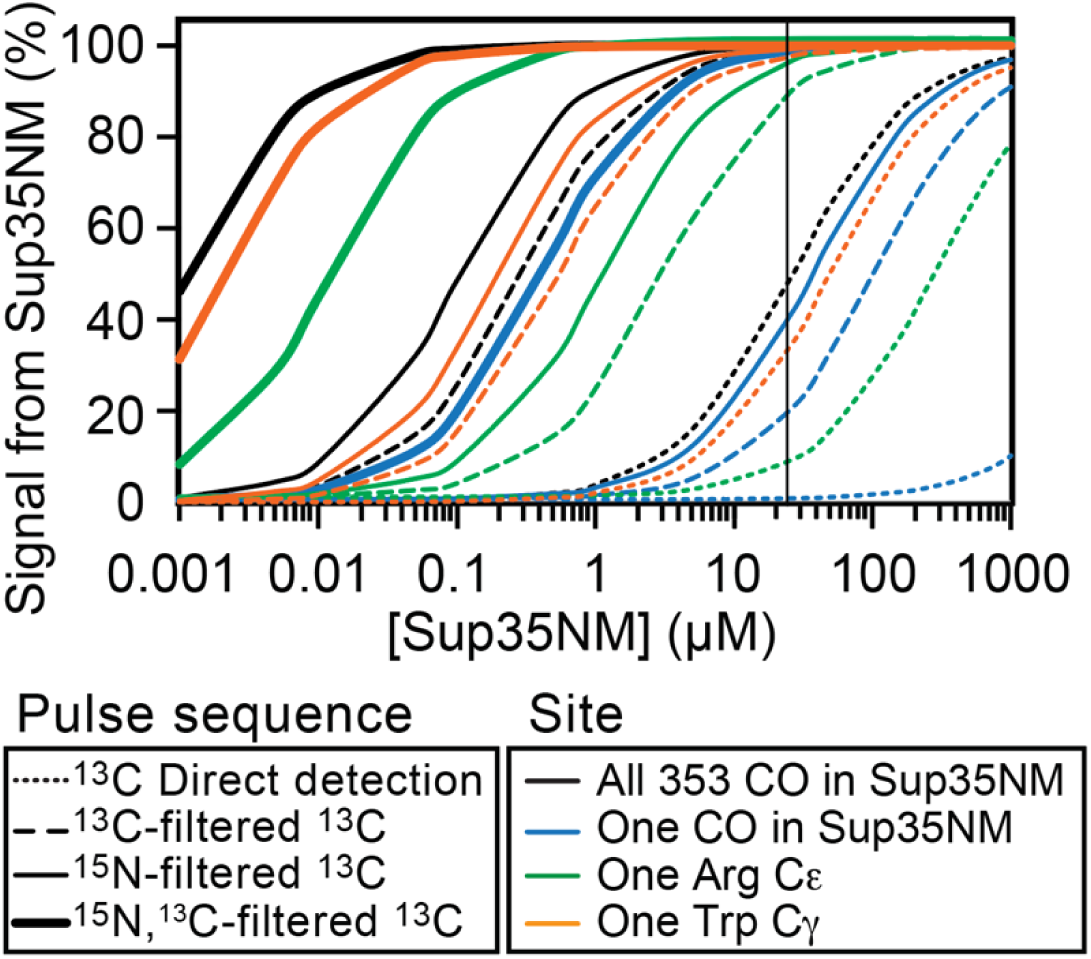
Specificity for the isotopically enriched protein of interest diluted in natural abundance cellular milieu depends upon the concentration of the protein of interest, the selectivity of the pulse sequence, and the degeneracy of the chemical shift of the site. The percent of the signal that is derived from isotopically enriched Sup35NM rather than from the natural molecules in the cellular milieu increases as the concentration of Sup35NM added to the sample increases. When Sup35NM is present at 25 µM (vertical line), the signal from all the carbonyl carbons in Sup35NM and from the cellular milieu is calculated to be of equal magnitude (dotted black line). Selection for sites with an adjacent isotopically labeled carbon (dashed lines) or nitrogen (solid lines) decreases the contribution from the cellular milieu by two orders of magnitude. Selection for a site through two adjacent isotopically labeled sites (thick lines) decreases the contribution from the cellular milieu by four orders of magnitude. All amino acids have at least one carbonyl moiety so specific detection of a single labeled CO site requires high concentrations and/or very selective pulse sequences (blue lines). Single sites in less abundant amino acids and/or in unique chemical moieties can be specifically detected at lower concentrations. About 5% of the amino acids in yeast are Arg (green) while only 1% are Trp (orange). With the use of highly selective pulse sequences these sites can be specifically detected (>98%) at 100 nM or lower concentrations.

We experimentally determined the selectivity ratio for a uniformly isotopically enriched Sup35NM at 25 µM diluted in lysates of yeast harboring the strong prion phenotype. To do so, we collected ^13^C detected proton-carbon cross-polarization (^13^C-CP) spectra(Pines, Gibby et al. 1973) of lysates of yeasts grown on isotopically naturally abundant media in the presence and absence of 25 µM of isotopically-enriched Sup35NM (Figure 4). Both spectra had peaks for carbonyl, aromatic, and aliphatic carbons, which are plentiful in proteins, and peaks for alcohol and ester carbons, which are plentiful in cell wall carbohydrates (∼70-106 ppm). The protein peaks were more intense in the spectra of isotopically enriched Sup35NM diluted in cellular lysates (Figure 4, green) than in the spectrum of lysates alone (Figure 4, blue), while the cell wall carbohydrate peak intensities were similar. Subtraction of the spectrum of natural abundance lysate alone from the spectrum of Sup35NM diluted in natural abundance lysates removes the spectral contribution from lysates and the resulting spectrum should report only on the Sup35NM present in the sample. The intensity of the carbonyl peak in the subtracted spectrum (Figure 4, yellow) was the same as the intensity of the carbonyl peak in the spectra of lysates alone (Figure 4, blue), in line with the theoretical selectivity ratio of one to one. The theoretically-determined and the experimentally-determined selectivity ratios for 25 µM isotopically enriched protein diluted in natural abundance lysates agree.

**Figure 4:**
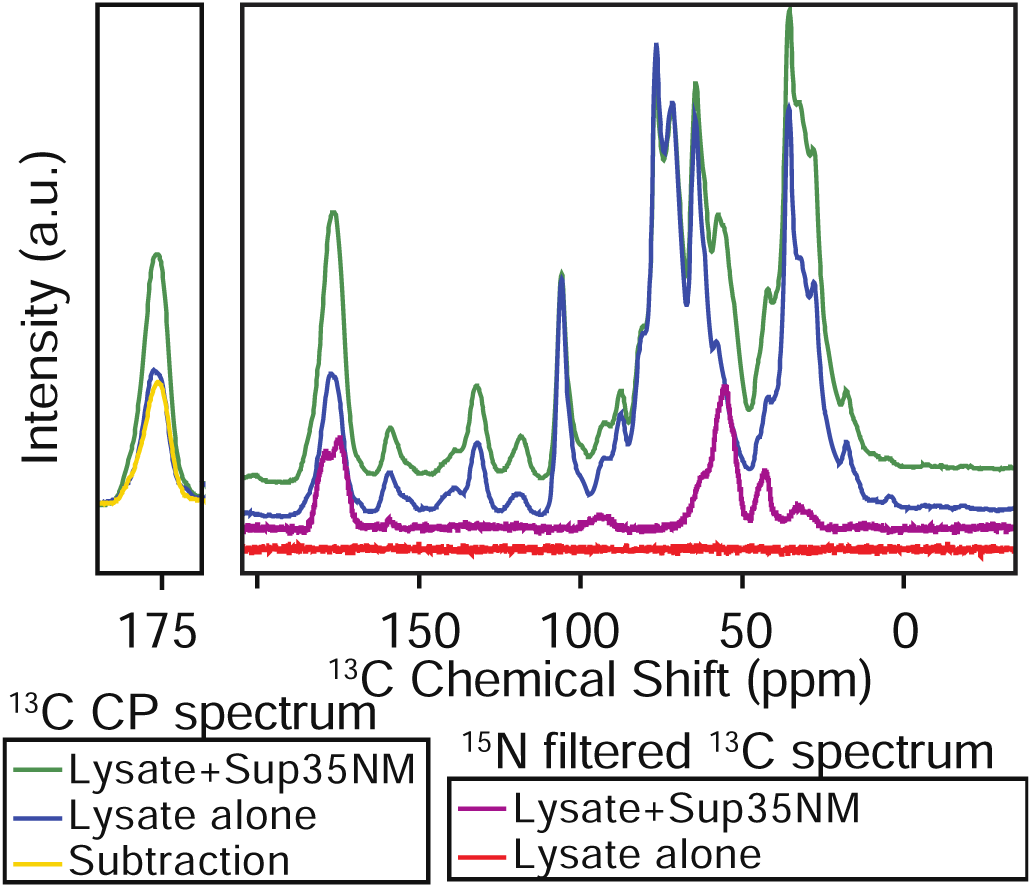
The protein of interest and naturally abundant cellular lysates have predictable magnitudes of contribution to the spectrum. The ^13^C spectra of natural abundance lysate alone (blue) has many signals while the ^15^N filtered ^13^C spectra has none (red). The ^13^C spectra of 25 μM of isotopically enriched Sup35NM diluted in natural abundance lysates (green) has many signals while the ^15^N filtered ^13^C spectra reports uniquely on the isotopically enriched Sup35NM (purple). Subtraction of the blue spectrum from the green spectrum results in the yellow spectrum (left). All data were collected at 600 MHz with 395 GHz microwave irradiation at 104 K with a recycling delay of 4s.

Experiments that report only on sites that are near other NMR-active nuclei – something that is probable in the isotopically enriched protein (100% of sites) and improbable in the cellular lysates - increase the selectivity of the experiment for the protein of interest. Considering only the ^13^C labeled carbonyl carbons that are directly bonded to another ^13^C site, the calculated contribution from a 30 kDa protein to the total signal is one hundred times larger than the contribution from the yeast lysate and considering only the ^13^C labeled carbonyl carbons that are directly bonded to another ^15^N site, the calculated contribution from the uniformly isotopically enriched protein to the total signal is two hundred times larger than the contribution from the yeast lysate. We experimentally assessed the selectivity ratio for a uniformly isotopically enriched Sup35NM at 25 µM diluted in yeast lysates. To do so, we collected ^15^N filtered ^13^C detected spectra (TEDOR) (Jaroniec, Filip et al. 2002) of lysates of yeasts grown on isotopically naturally abundant media in the presence and absence of 25 µM of isotopically-enriched Sup35NM (Figure 4)(Pines, Gibby et al. 1973). The spectrum of isotopically enriched Sup35NM diluted in cellular lysates had peaks for the carbonyl and Cα carbons, which are adjacent to the amide nitrogen in the protein backbone and, as expected, did not have peaks for aromatic and alcohol and ester carbons (∼70-106 ppm). Likewise, as expected at this signal to noise level, no carbon signals were detected in spectrum of lysates alone (Figure 4). Thus, using pulse sequences that select for adjacent isotopes, specific detection of uniformly isotopically enriched proteins in natural abundance yeast lysates at concentrations in the hundreds of nanomolar is theoretically possible (Figure 3).

### Specific detection of single protein sites in cellular milieu

The selectivity ratio for an isotopically enriched protein diluted in natural abundance lysates depends not only on the concentration of the protein but also the number of isotopically enriched sites in the protein of interest. Therefore, we determined the selectivity ratio in the limiting case where only single carbonyl carbon is uniformly isotopically enriched. Sup35NM has 253 amino acids and some of these amino acids have carbonyl carbons in the sidechain as well as the backbone so Sup35NM that is isotopically enriched only at a single site would have ∼353 times fewer labeled sites so the contribution of a single labeled amino acid is ∼353 times smaller. Thus, in a ^13^C NMR experiment that is directly-detected, a protein harboring a single isotopically labeled amino acid must be present at 6.25 mM and, in an ^15^N-filtered experiment, the protein must be present at 35 µM for the ^13^C carbonyl signal from a single labeled site to account for half of the ^13^C carbonyl signal (Figure 3, blue). However, the carbonyl carbon site represents the most challenging case for detection of a single labeled amino acid in a protein because the amino acids in the protein and the cellular milieu have an average of ∼1.25 carbonyl carbons and all carbonyl carbons fall within a relatively narrow range of chemical shifts. The selectivity ratios are better for sites and amino acids with distinct, rather than degenerate, chemical shifts. For example, arginine accounts for 5% of the amino acids in yeast and tryptophan accounts for 1% of the amino acids in yeast. Thus, detection of an arginine Cε site or a tryptophan Cγ site is 25- and 125- fold more selective, respectively, than detection of a carbonyl carbon because there are many fewer atoms in the sample with similar chemical shifts (Figure 3, green, orange). Use of pulse sequences that select for three adjacent atoms, instead of two, will be 100- fold more selective. For the CO, which is the most degenerate site, selection for three adjacent atoms will enable specific detection of a single labeled CO site at low micromolar concentrations (Figure 3, thick line). For a Trp site, selection for three adjacent atoms will enable specific detection for concentrations of tens of nanomolar in natural abundance cellular milieu. The selectivity ratio of uniformly isotopically enriched proteins diluted in natural abundance lysates depends upon both the concentration of the protein and number of isotopically enriched sites with degenerate chemical shifts. Thus, site-specific detection of single amino acid sites in a protein with distinct chemical shifts in cellular milieu for proteins at high nanomolar concentrations is theoretically possible.

Finally, isotopic depletion of the lysate background could potentially improve selectively for the isotopically enriched protein. We experimentally determined the selectivity ratio for a uniformly isotopically enriched Sup35NM at 25 µM diluted in lysates of yeast harboring the strong prion phenotype grown in isotopically depleted media. Because laboratory yeast strains are genetically altered to harbor nutritional defects, we grew yeast in complete synthetically-defined media (SD-CSM) which was made with ^13^C-depleted glucose (0.01%) and ^15^N-depleted ammonium sulfate (0.001%) as carbon and nitrogen sources but is supplemented with natural abundance amino acids and nutrients. We collected ^13^C detected ^13^C-CP spectra of lysates of yeasts grown on isotopically-depleted media in the presence and absence of 25 µM of uniformly isotopically-enriched Sup35NM (Figure S4A). The peak positions and relative intensities of the spectra of the isotopically depleted lysates alone were the same as those for natural abundance lysates alone but the total intensity of the isotopically-depleted lysates was half that of the natural abundance lysates (Figure S4B). Growth on isotopically depleted media resulted in a 50% reduction in the isotopic content of the lysates. This degree of depletion is in line with expectations because yeast will preferentially use the natural abundance amino acids present in the media. We next compared the spectra of isotopically depleted lysates in the presence and absence of Sup35NM and found that the signal intensities for the cell wall components in these spectra were similar (Figure S4A) while the signal intensities for proteins were four times more intense in the spectrum of the sample that contained Sup35NM. Subtraction of the spectrum of the isotopically-depleted lysates alone from the spectrum of Sup35NM in isotopically-depleted lysates removes the spectral contribution from lysates and the resulting spectrum reported only on the Sup35NM present in the sample (Figure S4A, orange). The intensity of the subtracted spectrum (Figure S4A, gold) is twice that of the depleted lysate alone (Figure S4A, light blue), in agreement with the degree of isotopic depletion of the lysates. Thus, growth of cells on isotopically-depleted nutritional sources decreased the signal contribution from the cellular background proportional to the degree of isotopic depletion. If the isotope content in the yeast lysates is reduced further, which could be accomplished by growing prototrophic yeast on minimal media made with isotopically depleted carbon and nitrogen sources, specific detection of uniformly isotopically enriched proteins in isotopically depleted yeast lysates at concentrations in the low nanomolar range is theoretically possible(Costello, Xiao et al. 2019). Thus, for samples with homogenously dispersed AMUPol and well-ordered protein sites, the concentration of proteins that can be specifically detected in cellular lysates far surpasses the practical limit of experimental sensitivity.

Finally, to assess the site-specific detection of single amino acid sites in a protein in cellular milieu, we collected two-dimensional carbon-nitrogen correlation spectra (zTEDOR) of 25 µM Sup35NM in yeast lysates. The DNP enhancement was 72 for this sample, which was made with fully protonated buffer and yeast lysates and cryoprotected with 15% glycerol. The signal to noise ratio for the carbonyl carbon peak was 115 after 23 hours of acquisition time. The resolution of this spectrum enabled identification of amino acids with distinct ^13^C-^15^N chemical shifts (Figure 5). Of the six amino acids with adjacent nitrogen-carbon pairs with chemical shifts that are distinguished by 10 ppm or more from the average for protein backbone chemical shifts, Sup35NM contains all but tryptophan (Figure 5A) with 25 lysines, 23 glycines, 14 prolines, 2 arginines and 1 histidine. There were resolvable peaks in the spectra for the lysine C^ε^-N^ζ^, the glycine C^α^-N, the proline C^α^-N & C^δ^-N, and the arginine C^δ^-N^ε^ (Figure 5A). These peaks had signal to noise ratios that ranged from 150:1 for the lysine peak to 18:1 for the arginine peak (Table S1). Because peak intensities for different chemical moieties are not necessarily quantitative, we collected a 2D zTEDOR spectra of purified uniformly isotopically enriched Sup35NM that had been polymerized into the amyloid form using lysates from strong [*PSI^+^*] cells as a template(Frederick, Debelouchina et al. 2014). The relative ratios of the peak heights and SNR for lysine, glycine, proline and arginine are statistically indistinguishable for Sup35NM diluted into lysates and purified Sup35NM (p > 0.42, student’s t.test, paired). Peaks for neither histidine, which is present in Sup35NM but likely has multiple protonation states under these conditions, nor tryptophan, which is not present in Sup35NM but is a component of the lysates, were detected in either the two-dimensional spectra or the projections of the relevant regions (Figure 5). Thus, site-specific detection of resolved amino acids sites for an isotopically enriched protein at low concentrations diluted in fully-protonated natural abundance cellular milieu is possible in tractable acquisition times.

**Figure 5.**
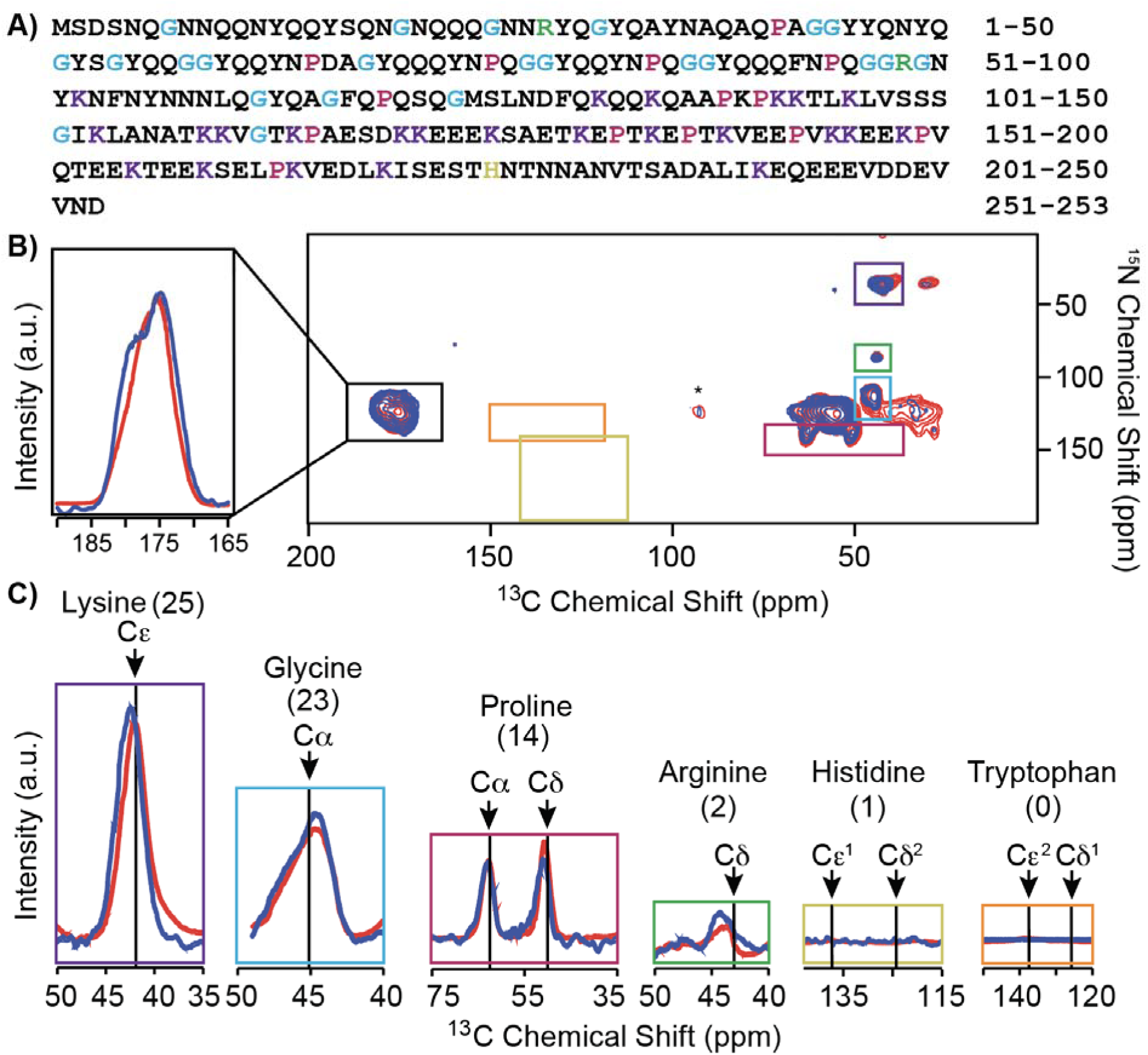
Specific detection of isolated sites in Sup35NM in cellular milieu. (A) Protein sequence of Sup35NM. Lysines (purple), glycines (blue), prolines (pink), arginines (green) and histadine (yellow) are highlighted. (B) Two-dimensional ^15^N-^13^C correlation spectra (zTEDOR). Uniformly labeled Sup35NM in protonated, natural abundance strong [PSI^+^] yeast lysate (blue) with 85% protonated buffer and 15% d_8_-glycerol with 5 mM AMUPol, n=1280, acquisition time = 23 hours, ε=72. Purified amyloid fibers (red) with 10% protonation, 60% ^13^C depleted d_8_-glycerol with 10 mM AMUPol, n=128, acquisition time = 2.5 hours ε=44. Data were collected at 600 MHz with 12.5 kHz MAS at 104 K with a recycle of 4 seconds. Asterisks indicate spinning side bands. (C) Projections of regions of the zTEDOR for amino acids with unique chemical shifts for the lysate sample (colors, projected regions marked in (B) by boxes) overlayed on the purified fibril spectra (blue). The number of each amino acid in Sup35NM is indicated in paratheses. Arrows indicate the average chemical shifts from the BMRB for each site. Signal to noise ratios, integrated peak areas and centers are reported in Table S1.

To assess the influence of cellular milieu on the conformation of Sup35NM, we compared the spectra of Sup35NM that had been polymerized into the amyloid form using lysates from strong [*PSI^+^*] cells as a template in the presence and absence of cellular milieu. In both samples, the average chemical shifts for the carbonyl, C^α^ and backbone nitrogen sites for all sites were consistent with β-sheet conformations, with larger shifts towards β-sheet chemical shift values for the Sup35NM assembled in cellular milieu (Table S1). The chemical shifts in both spectra for sites in the sidechains all fall within a standard deviation of the average value reported in the BMRB, but differ from each other; the center of the lysine C^ε^ and arginine C^δ^ peaks were larger by 0.5 ppm when Sup35NM was polymerized in the presence, rather than the absence, of cellular milieu. The peak widths of the lysine, glycine and carbonyl carbon peaks were all broader for Sup35NM when polymerized in the presence rather than the absence of cellular milieu, reflecting a wider range of sampled conformations in the cellular milieu. Indeed, the carbonyl carbon peak for Sup35NM assembled in cellular milieu had a pronounced shoulder at chemical shift values consistent with the carbonyls in asparagine and glutamine sidechains, indicating that the sidechains are more ordered in the presence of cellular milieu than in its absence (Figure 5). This indicates that the cellular milieu can, and does, influence protein conformations.

## DISCUSSION

The sensitivity enhancement from DNP NMR is theoretically sufficient to detect endogenous concentrations of isotopically-enriched proteins in biological environments. Yet, even with DNP, experiments are still sensitivity limited. To improve experimental sensitivity, we optimized sample composition and found that moderate concentrations of cryoprotectants improve absolute sensitivity without compromising enhancements and that per-deuteration does not improve DNP performance. The latter result indicates that good DNP performance can be expected in both per-deuterated and fully protonated cellular systems. Because at low target protein concentrations, the amount NMR-active isotopes in cellular biomass from natural abundance could be larger than that in the target protein, we experimentally validated theoretical calculations of the limit of specificity for an isotopically enriched protein in natural abundance cellular milieu. We establish that, using pulse sequences that are selective for adjacent NMR-active nuclei, it is possible that proteins can be specifically detected in cellular milieu at concentrations in the hundreds of nanomolar. In a fully protonated sample, cross peaks for isolated sites can be detected in under a day of acquisition time for the yeast prion protein, Sup35NM, when it is present at a concentration of 25 µM in concentrated yeast lysates. We found while there are no intensity differences in the spectra of purified Sup35NM and Sup35NM diluted in cellular lysates, which indicates that the cellular lysates did not contribute to the spectra, there were difference in the chemical shifts, which indicates that the cellular milieu can and does influence protein conformations.

Because cellular samples are sensitivity limited, we investigated the effect that the amount of cryoprotectant and level of deuteration had on DNP performance. Here we found that a cryoprotectant concentration of 15% (*v/v*) glycerol resulted in the highest absolute sensitivity for samples of proteins diluted in cellular milieu because samples with lower concentrations of cryoprotectants had higher DNP enhancements as well as higher fill factors. While in this work, we limited our investigations to glycerol because isotopically depleted versions are commercially available, this is likely extendable to moderate concentrations of other cryoprotectants for cellular milieu (Xiao, Ghosh et al. 2021). For insoluble biomass, lowering the concentration of the cryoprotectant decreased the homogeneity of the dispersion of the polarization agent throughout the sample. This contrasts with reported results in mammalian cell systems (Ghosh, Xiao et al. 2021, Xiao, Ghosh et al. 2021) as well as with results reported here with whole yeast lysates; the homogeneity of samples of mammalian cell derived milieu cryoprotected with 15% glycerol remained high as did whole lysates of yeast cells. Moreover, we found that although deuteration levels have a modest effect on the DNP properties of AMUPol, the deuteration level of the buffer and the cellular components did not alter the DNP properties of AMUPol in biological milieu. This indicates that the deuteration level doesn’t significantly affect the DNP performance for these complex samples and, more generally, indicates that with modern radicals, per-deuteration may not be required at all for good DNP performance. Thus, here we find that in complex biological settings, reducing the amount of added cryoprotectant doubles the absolute sensitivity and that per-deuteration, which can alter the behavior of biological and cellular systems, is not a requirement for optimal DNP performance. Optimized sample compositions combined with modern polarization agents (Lund, Casano et al. 2020, Stevanato, Casano et al. 2020, Harrabi, Halbritter et al. 2022, Yao, Beriashvili et al. 2022) will enable detection of proteins at concentrations in the high nanomolar range in tractable acquisition times.

The endogenous concentrations of many proteins range from hundreds of micromolar to tens of nanomolar. For samples that are homogenously doped with the polarization agent, the specificity limit for an isotopically enriched site in natural abundance cellular milieu depends upon the abundance of sites with chemical shifts that are degenerate with that of the isotopically-enriched site. Using pulse sequences that select for adjacent isotopically enriched sites, the theoretical specificity limit can range from hundreds of micromolar for a carbonyl site to single digit micromolar for sites with well-resolved chemical shifts. The specificity limit can be lowered by further isotopic depletion of the cellular background (Costello, Xiao et al. 2019). However, the specificity limit can also be lowered by an additional two orders of magnitude by using more selective pulse sequences, such as those that incorporate selection steps that require not just two, but three, adjacent isotopically enriched sites(Pauli, Baldus et al. 2001, Heise, Seidel et al. 2005). Finally, while experimental specificity is an important consideration for samples that are homogenously doped with polarization agents where the analyte is diluted in an environment containing concentrated chemically similar natural abundance molecules, covalent attachment of the polarization to the target would effectively eliminate the specificity limit. (Viennet, Viegas et al. 2016, Lim, Ackermann et al. 2020) Likewise, experimental specificity is not a concern for molecules with chemical shifts that are distinct from those of the components of the cellular biomass(Schlagnitweit, Sandoz et al. 2019, Bertarello, Berruyer et al. 2022). Overall, with the currently available suite of DNP polarization agents, the specificity limit covers the range of endogenous concentrations for many proteins and is far lower than the practical sensitivity limit.

Here, we determined that by using selective pulse sequences, well-resolved single sites can potentially be specifically detected in yeast lysates when present at high nanomolar concentrations. Such calculations can be useful for other cellular systems. While not yet experimentally validated, the theoretical specificity limit for a single carbonyl site in mammalian cells is an order of magnitude lower than that for yeast because the protein density of the mammalian cells is lower than that of yeast (Milo 2013) and the relative abundance of the amino acids differs (Dietmair, Hodson et al. 2012, Oftadeh, Salvy et al. 2021). For the same reasons, the specificity in bacterial cells is expected to be an order of magnitude higher. Understanding the specificity limitations is useful for the design and interpretation of experiments of proteins in biological settings. For example, there are only two arginines in Sup35NM and the arginine C^δ^-N^ε^ correlations were detected with a SNR of 18:1 after 23 hours of signal averaging for 25 Sup35NM at 25 µM. However, for this protein system, the broader line widths under DNP conditions and chemical shift degeneracy precludes observation of single sites. Specific and segmental isotopic labeling effectively reduce chemical shift degeneracy of this protein (Frederick, Michaelis et al. 2017, Ghosh, Dong et al. 2018). Understanding the specificity limit defines the sample composition and choice of pulse sequences best suited to specifically detect unique sites for proteins at physiological concentrations from natural abundance isotopes present in the cellular milieu.

Prior work on the Sup35NM system indicated that a region that was intrinsically disordered in purified samples of Sup35NM fibrils underwent a large change in secondary structure when the protein was assembled into the amyloid form in cellular lysates (Frederick, Michaelis et al. 2015). In that work, the uniformly isotopically enriched Sup35NM at a concentration of 1 µM was diluted in isotopically depleted yeast lysates. The resulting increase in Sup35NM concentration in that sample was within range of endogenous levels since Sup35 is normally present at concentrations of 2.5 to 5 µM (Ghaemmaghami, Huh et al. 2003). In this work, the Sup35NM was present at a concentration of 25 µM, altering the stoichiometries of this protein with potential interactors by an order of magnitude, a perturbation that may potentially titrate out some interactions. Prior work reported on ^13^C-^13^C correlations, particularly those for well-resolved sites like the backbone C’-C^α^, the lysine C^δ^-C^ε^ and the proline C^γ^-C^δ^. The present work reports on ^13^C-^15^N correlations. The sites that are most comparable to prior work are the backbone C’-N (which, in this work, includes contributions from the C’-N in the sidechains of Asn and Glu), the lysine C^ε^-N^ζ^, and the proline C^δ^-N. The increase in β-sheet content for Sup35NM assembled in lysates relative to purified settings is maintained in both samples. However, while the chemical shifts of the sites in the lysine and proline sidechains differ for fibrils assembled in lysates and purified settings regardless of Sup35NM concentration, the magnitude of these differences, particularly for the proline C^δ^, is smaller in the sample made with non-endogenous concentrations of Sup35NM. Because the changes in chemical shift are more modest for sites in the region of intrinsic disorder when Sup35NM is present at 25 µM, this suggests that the conformational rearrangement seen at lower concentrations is a result of specific interactions, rather than crowding. This further suggests that the relative stoichiometry of the protein of interest to the components of the cellular environment is an important experimental consideration, particularly for proteins that make specific interactions with cellular constituents.

## MATERIALS AND METHODS

### Sample preparation

Sup35NM was expressed and purified as described elsewhere(Serio et al., 1999). Uniformly-labeled ^13^C ^15^N Sup35NM samples were prepared by growing BL21(DES)-Rosetta *Escherichia coli* in the presence of M9 media with 2 g L^-1^ D-glucose ^1^H,^13^C_6_ and 1 g L^-1^ of ^15^N ammonium chloride (Cambridge Isotope Labs, Cambridge, MA). Purified, lysate-templated NM fibrils samples for the purified fiber sample were prepared as described elsewhere(Frederick et al., 2014)(Frederick, Michaelis et al. 2015), except that 5 mM AMUPol was used as the polarization agent. (Sauvée, Rosay et al. 2013)

### Cell lysate samples for DNP

Cell lysate samples were prepared as previously described. (Frederick, Michaelis et al. 2015) (Costello, Xiao et al. 2019), with minor modifications. Briefly, phenotypically strong [*PSI^+^*] yeast were grown in a 50 mL culture volume at 30 °C to mid-log phase in SD-CSM media made with protonated carbon sources and either 100% H_2_O or 100% D_2_O. Because we use protonated carbon sources, the final deuteration level for the lysates was 70% as determined by solution state NMR, as expected(Leiting et al., 1998). Isotopically depleted yeast lysates were made from yeast grown on SD-CSM made with yeast nitril base without amino acids and without ammonium sulfate (BD Scientific) supplemented by ^13^C depleted D-Glucose (99.9% Cambridge Isotopes) and ^15^N depleted ammonium sulfate (Sigma). Cells were collected by centrifugation (5 min, 4000 x g) and washed once with water or D_2_O, depending upon final level of sample perdeuteration. Pellets were suspended in 200 µL of lysis buffer (50 mM Tris-HCl pH 7.4, 200 mM NaCl, 2 mM TCEP, 5% *d_8_*-^13^C depleted glycerol, 1 mM EDTA, 5 ug/mL of aprotinin and leupeptin and 100 µg/mL Roche protease inhibitor cocktail. Deuteration of the lysis buffer was adjusted as required.) Cells were lysed by bead beating with 500 µm acid washed glass beads for 8 minutes at 4 °C. After bead beating, the bottom of the Eppendorf tube was punctured with a 22G needle and the entire lysate mixture was transferred to a new tube. Purified denatured ^13^C^15^N-labeled Sup35NM was diluted at least 150-fold out of 6 M GdHCl to a final concentration of 5 µM and the mixture was allowed to polymerize, quiescent, at 4 °C for 24 to 48 hours. The insoluble portion of the sample was collected by centrifugation at 20000 x g for 1 hour at 4 °C and removal of the supernatant. The ∼30 µL pellet was resuspended in various amounts of 100% *d_8_*-^13^C depleted glycerol and transferred to a 3.2 mm sapphire rotor. The final radical concentration was 5 mM of AMUPol (Sauvée, Rosay et al. 2013). The isotopically depleted cell lysate samples were made analogously, except that yeast cells were grown in SD-CSM media made with 2% (w/v) protonated ^13^C-depleted glucose (99.9% ^12^C, *Cambridge Isotope Labs*) as the carbon source. Whole yeast lysate samples were made by growing the w303 strain of *S. cerevisiae* that were corrected for auxotrophies at all loci were cultured in synthetic defined yeast media (SD-min) made with 2 % (w/v) D-glucose-^13^C_6_ (Sigma), 0.225 % (w/v) ^15^NH_4_Cl (Sigma), and 0.17% nitrogen base without ammonium sulfate and amino acids (Difco). Cells were grown at 30 °C to mid log phase, collected by centrifugation and washed twice with D_2_O. Cells were buffered with a phosphate buffer (10 mM potassium phosphate, 100 mM NaCl, pH = 6.0) and suspended in *d_8_*-^12^C-glycerol/D_2_O/H_2_O for a final composition of 15:75:10 with 10 mM AMUPol. The cells from the suspension were pelleted into a 3.2 mm rotor and lysed freeze-thawing the yeast cells in the rotor three-times. The yeast lysate was stored at - 80°C until data collection

### Immunohistochemistry

Cell lysate samples were made as described above. Sup35NM was visualized using an antibody raised against residue 125-253 of the protein(GenScript). Cell lysates were fractionated by SDS-PAGE, transferred to nitrocellulose and probed with the anti-Sup35NM antibodies. Lysate samples were denatured by incubation at 95 °C for 10 minutes in the presence of 2% SDS before fractionation to denature amyloid aggregates. Secondary antibodies were coupled to horseradish peroxidase. Blots were visualized by a standard ECL analysis.

### Spectroscopy

Dynamic nuclear polarization magic angle spinning nuclear magnetic resonance (DNP MAS NMR) experiments were performed on 600 MHz Bruker Ascend DNP NMR spectrometers with 7.2 T Cryogen-free gyrotron magnet (Bruker), equipped with a ^1^H, ^13^C, ^15^N triple-resonance, 3.2 mm low temperature (LT) DNP MAS NMR Bruker probe (600 MHz). The sample temperature was 104 K, at MAS frequency of 12 kHz. The DNP enhancements on these instrumentation set-ups for a standard sample of 1.5 mg of uniformly ^13^C, ^15^N labeled proline (Isotech) suspended in 25 mg of 60:30:10 *d_8_*-glycerol:D_2_O:H_2_O containing 10 mM AMUPol varied between 80 and 140. The DNP enhancements that were measured close in time on the same spectrometer were compared quantitively within the set and qualitatively between sets.

In ^13^C cross-polarization (CP) MAS experiments, the ^13^C radio frequency (RF) amplitude was linearly swept from 75 kHz to 37.5 kHz with an average of 56.25 kHz. ^1^H RF amplitude was kept at 68∼72 kHz for CP, 83 kHz for 90 degree pulse, and 85 kHz for ^1^H TPPM decoupling with phase alternation of ± 15° during acquisition of ^13^C signal. The DNP enhancements were determined by comparing 1D ^13^C CP spectra collected with and without microwaves irradiation. For *T_B,on_* measurements, recycle delays ranged from 0.1 s to 300 s. To determine the *T_B,on_*, the dependence of the recycle delay on both ^13^C peak intensity or volume was fit to the equation 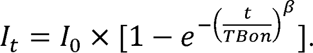 For ^13^C-^15^N 1D and 2D correlations, a 1.92 ms TEDOR sequence was applied with ^13^C and ^15^N pulse trains at 55.5 kHz and 41.7 kHz, respectively. A total of 64 complex *t_1_* points with an increment of 80 μs were recorded. The recycle delay was 4.0 s and the same ^1^H decoupling was applied. Data were processed using NMRPipe software and analyzed in Sparky.

## SUPPLEMENTAL INFORMATION

Supplemental information is available for this article.

## ACKNOWLEDGEMENTS

W.N.C. was supported by a graduate research fellowship from the NSF and NIH MB T32 GM008297. This work was supported by grants from the National Science Foundation [1751174]; the Welch Foundation [1-1923-20200401]; the Lupe Murchison Foundation, and the Kinship Foundation (Searle Scholars Program) to K.K.F. The National High Magnetic Field Laboratory is supported by the NSF (DMR1644779 and DMR-2128556) and by the State of Florida. The 14.1 T DNP system at NHMFL is funded in part by NIH S10 OD018519 (magnet and console), NSF CHE-1229170 (gyrotron), and NIH P41 GM122698.

## Notes

### Competing Interest Statement

The authors have declared no competing interest.

### Summary of Updates

Improve readability and interpretability of figures.

