## Supplemental Information for "DNP-assisted solid-state NMR enables detection of proteins at nanomolar concentrations in fully protonated cellular environments"

Current affiliation:

4. National Institute of Chemistry, Hajdrihova 19, 1001 Ljubljana, Slovenia

*To whom correspondance should be addressed: Kendra K. Frederick

https://orgic.org/0000-0001-6731-8978

https://orcid.org/0000-0002-4548-4833

https://orcid.org/0000-0002-3570-9787

https://orcid.org/0000-0001-6472-8429

https://orcid.org/0000-0002-1656-5167

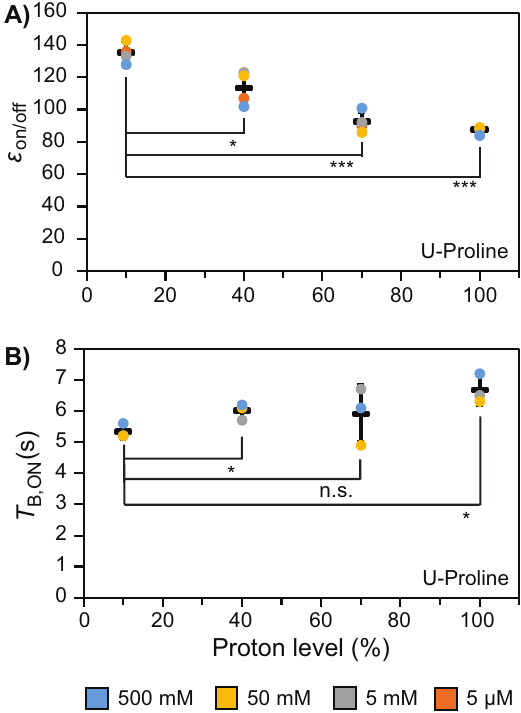

***Figure S1****:* *DNP performance decreases ~30% in proline standard samples with high protonation levels. DNP performance is independent of analyte concentration. (A)  DNP enhancement (ε_on/off_) and (B) Build-up times, T_B,ON_, from mono-exponential fits increase with increasing protonation levels of proline standard samples at 500 mM (blue), 50 mM (yellow), 5 mM (grey) and 5 µM (orange) measured by the ratio of on/off microwaves signal of the ^13^C-CP signal at the carbonyl decreases with increasing protonation levels. Data averages are shown in black. Standard deviation is marked by error bars.*

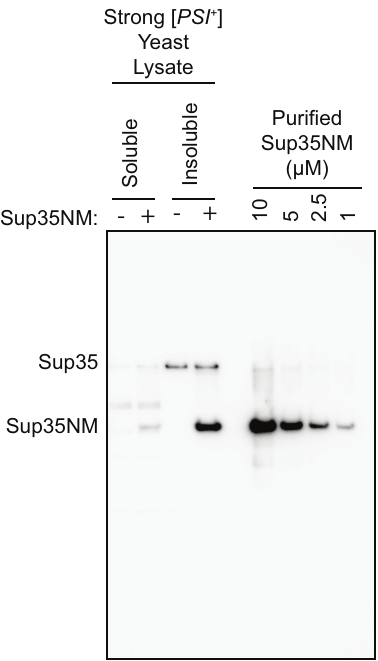

***Figure S2:*** *Sup35NM is present at 25 μM in DNP NMR cellular lysate samples.* *Semi-quantitative Western blot of DNP NMR samples with and without exogenously added Sup35NM at 5 μM in 200 μL lysis mixture of lysis buffer and 50 mL of strong [PSI+] yeast harvested at OD = 0.7. For DNP NMR samples the insoluble portion is harvested and concentrated into a ~30 μL pellet, making the final concentration 7.2x of that shown in the Western blot (~3.5 μM).*

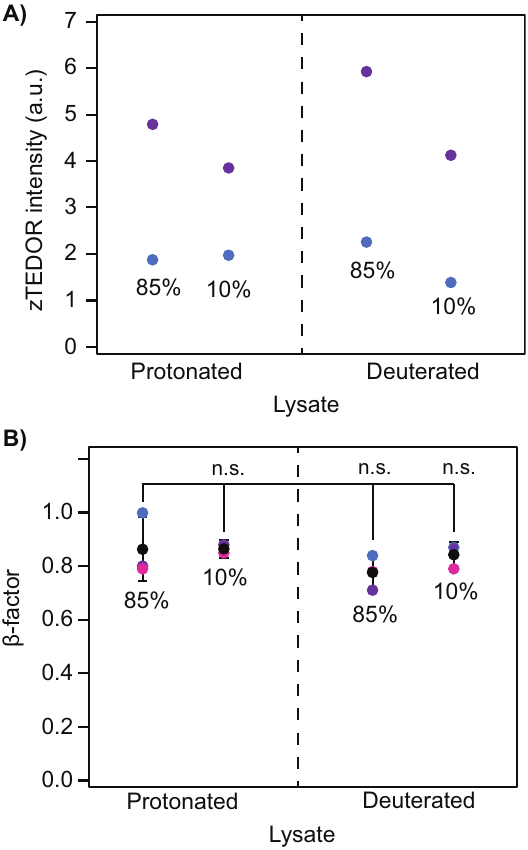

***Figure S3.*** *Protein specific signal and β-factor are not affected by protonation level.* ***A)*** *zTEDOR signal intensity of cellular milieu samples measured at the carbonyl.* ***B)*** *β-factor from fitting build-up times to a stretched exponential measured using integration of the carbonyl peak. All sample sets contained 15% d_8_-glycerol and 5 mM AMUPol biradical.*

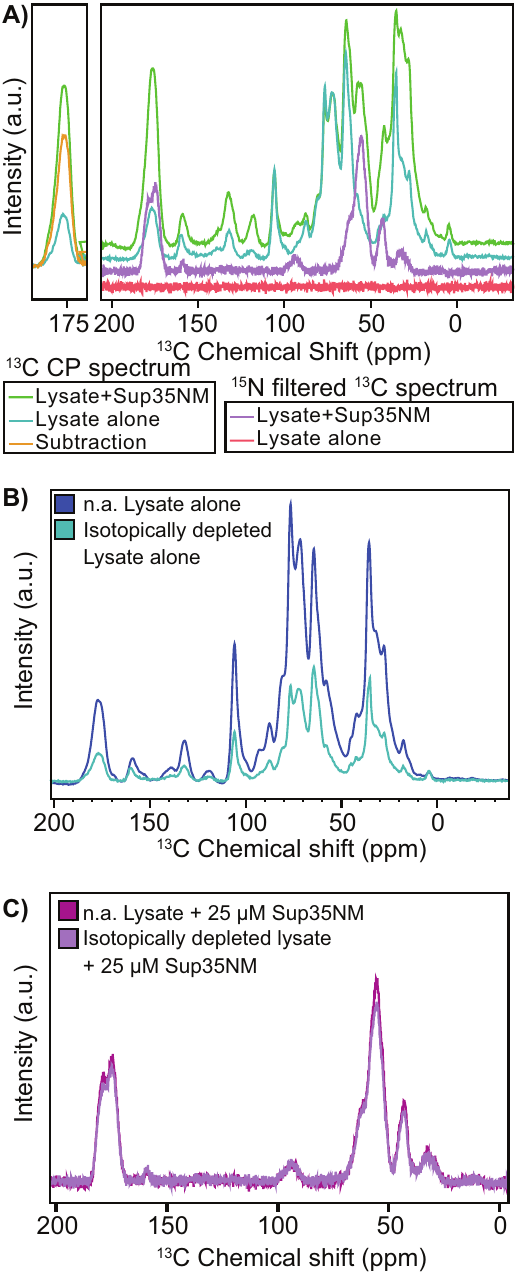

**Figure S4** Depletion of the cellular lysate background does not affect signal from the protein of interest. *The ^13^C spectra of isotopically depleted lysates alone (light blue) has many signals while the ^15^N filtered ^13^C spectra has none (red). The ^13^C spectra of 25 μM of isotopically enriched Sup35NM diluted in isotopically depleted lysates (green) has many signals while the ^15^N filtered ^13^C spectra reports uniquely on the isotopically enriched Sup35NM (purple). Subtraction of the blue spectrum from the green spectrum results in the orange spectrum (left). All data were collected at 600 MHz with 395 GHz microwave radiation at 104 K with a recycle delay of 4s. (B) Comparison of yeast lysates grown on natural abundance and isotopically depleted media. (C) Comparison of the ^15^N filtered ^13^C spectra of Sup35NM at 25 µM in natural abundance and isotopically depleted lysates.*

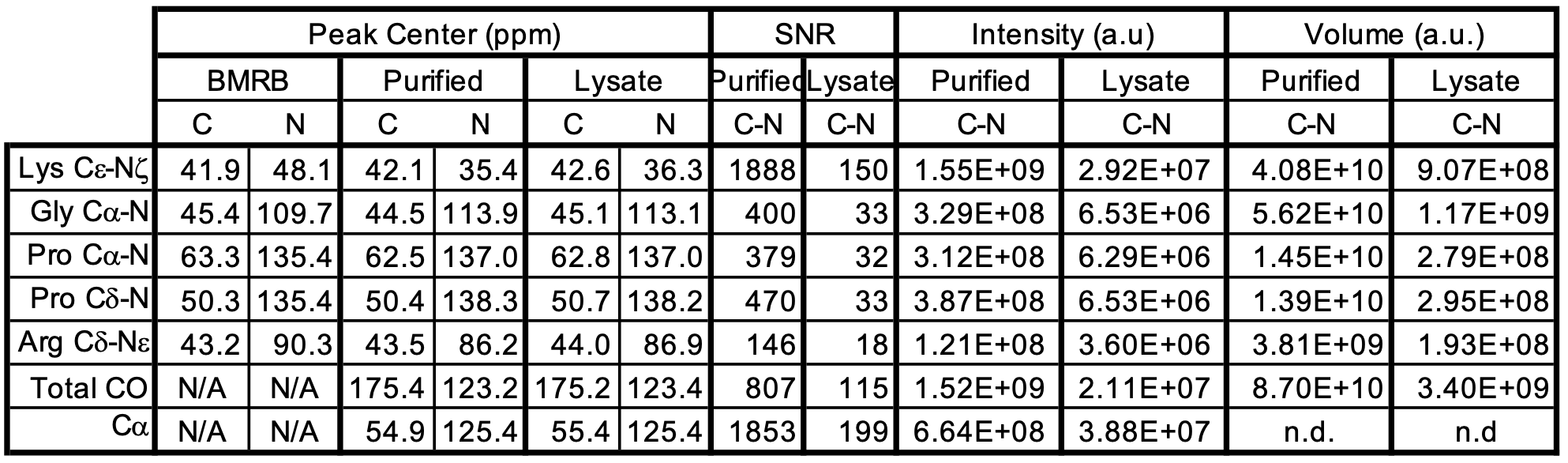

***Table S1****: Peak centers, signal-to-noise ratio (SNR), peak intensity and integrated volumes of from the 1D projections (normalized for the size of the projected range in the 15N dimension for unique amino acids in reported of the 30 kDa protein Sup35NM when it was templated into the amyloid form in buffer as when it was templated into the amyloid form in cellular lysates at 25 μM.*

***Calculation of the experimental specificity***

Carbon and Nitrogen specificity ratios were calculated in similar manners to reflect the abundance respectively of each in the protein component of yeast cells.

**Isotopic contribution from the insoluble protein carbonyl portion of the lysate**

Starting with 50 mL of yeast culture with an OD_600_ = 0.7 yields

50 mL * 0.7 OD_600_ * 10^7^ cells/(mL* OD_600_) = 3.5*10^8^ cells. (Sherman 1991, Groves, Falson et al. 1996)

The dry mass of a single yeast cell is 47.65 ± 1.05 pg (Łabędź, Wańczyk et al. 2017). Therefore, the estimated dry mass of the 50 mL yeast culture sample is

47.65*10^-9^ mg/cell * 3.5*10^8^ cells = 16.68 mg of dry mass.

About 40% of yeast biomass is composed of proteins(Van Hoek, Van Dijken et al. 1998, Gombert, Moreira dos Santos et al. 2001, Lange and Heijnen 2001, Oftadeh, Salvy et al. 2021).

16.68 mg * 40% = 6.67 mg of protein.

Carbonyl carbons compose a fraction of this mass. Using the percent mol contribution of each amino acid in yeast (Oftadeh, Salvy et al. 2021), we calculated the percentage of total carbonyls in the protein component of yeast to obtain the carbonyl weighing factor. Weighing factors for other sites can be calculated analogously.

| Amino acid | # of Carbons | Carbon weight (Da) | M.W. (Da) | Weighing factors | % mol | % mol * M.W.  (g) | % dry weight (mg) | Carbonyl dry weight (%) |
| --- | --- | --- | --- | --- | --- | --- | --- | --- |
| R | 1 | 12 | 174.2 | 0.07 | 3.86 | 672.41 | 5.31 | 0.37 |
| H | 1 | 12 | 155.2 | 0.08 | 1.93 | 299.54 | 2.37 | 0.18 |
| K | 1 | 12 | 147.2 | 0.08 | 6.57 | 967.10 | 7.64 | 0.62 |
| D/N* | 2 | 24 | 132.6 | 0.18 | 9.28 | 1230.53 | 9.72 | 1.76 |
| E/Q* | 2 | 24 | 146.7 | 0.16 | 15.48 | 2270.14 | 17.92 | 2.93 |
| S | 1 | 12 | 105.1 | 0.11 | 5.33 | 560.18 | 4.42 | 0.51 |
| T | 1 | 12 | 119.1 | 0.10 | 5.57 | 663.39 | 5.24 | 0.53 |
| C | 1 | 12 | 121.2 | 0.10 | 0.14 | 16.97 | 0.13 | 0.01 |
| G | 1 | 12 | 75.1 | 0.16 | 8.89 | 667.64 | 5.27 | 0.84 |
| P | 1 | 12 | 115.1 | 0.10 | 4.22 | 485.72 | 3.84 | 0.40 |
| A | 1 | 12 | 89.1 | 0.13 | 9.77 | 870.51 | 6.87 | 0.93 |
| V | 1 | 12 | 117.1 | 0.10 | 7.33 | 858.34 | 6.78 | 0.69 |
| I | 1 | 12 | 131.2 | 0.09 | 5.89 | 772.77 | 6.10 | 0.56 |
| L | 1 | 12 | 131.2 | 0.09 | 8.01 | 1050.91 | 8.30 | 0.76 |
| M | 1 | 12 | 149.2 | 0.08 | 1.14 | 170.09 | 1.34 | 0.11 |
| F | 1 | 12 | 165.2 | 0.07 | 3.76 | 621.15 | 4.90 | 0.36 |
| Y | 1 | 12 | 181.2 | 0.07 | 1.96 | 355.15 | 2.80 | 0.19 |
| W | 1 | 12 | 204.2 | 0.06 | 0.65 | 132.73 | 1.05 | 0.06 |
|  |  |  |  | Total | 99.78 | 12665.27 | 100.00 | 11.80 |

*D and N are Glx, E and Q are Asx, all of them have 2 COs

To determine how much carbonyl signal from natural abundance comes from the insoluble yeast lysate, we multiplied the protein dry weight by the natural abundance of ^13^C, which is 1.1%, and by the carbonyl weighing factor. The detectable ^13^C in protein carbonyls for a 50 mL yeast culture at OD_600_ = 0.7 is

6.67 mg of protein * 1.1% (^13^C natural abundance) * 11.8% carbonyl = 0.0087 mg = 8.7 µg.

These sample use only the insoluble portion of the cellular biomass and 66% of protein content remains in the insoluble portion for samples prepared in this way(Frederick, Michaelis et al. 2015). Therefore, there are:

8.7 µg of ^13^C carbonyl carbons from natural abundance * 66% = 5.7 µg

of detectable ^13^C in protein carbonyls in the insoluble natural abundance yeast lysate of the DNP NMR samples in this study.

**Isotopic contribution from the insoluble protein carbonyl portion of Sup35NM**

For the protein of interest, we calculated the number ^13^C labeled carbonyl carbons present in the uniformly labeled Sup35NM. Using the amino acid composition and the number of carbonyls in each amino acid we determined the formula weight of the carbonyls in the Sup35NM protein by taking the number of amino acid residues in Sup35NM multiplied by the number of carbonyls in each amino acid multiplied by the weight of ^13^C (13 Da) and we calculated the weighing factor for the carbonyl in the Sup35NM molecule. We divided the total carbonyl formula weight by the Sup35NM molecular weight of the isotopically [^13^C,^15^N]-labeled molecules to obtain the carbonyl weighing factor. Weighing factors for different labeling schemes can be obtained analogously.

| Amino Acid | # of residues in Sup35NM | # of carbonyl | Carbonyl F.W. in 13C,15N-Sup35NM |
| --- | --- | --- | --- |
| A | 15 | 1 | 195 |
| R | 2 | 1 | 26 |
| N | 27 | 2 | 702 |
| D | 9 | 2 | 234 |
| C | 0 | 1 | 0 |
| Q | 41 | 2 | 1066 |
| E | 23 | 2 | 598 |
| G | 23 | 1 | 299 |
| H | 1 | 1 | 13 |
| I | 3 | 1 | 39 |
| L | 8 | 1 | 104 |
| K | 25 | 1 | 325 |
| M | 2 | 1 | 26 |
| F | 4 | 1 | 52 |
| P | 14 | 1 | 182 |
| S | 15 | 1 | 195 |
| T | 11 | 1 | 143 |
| W | 0 | 1 | 0 |
| Y | 20 | 1 | 260 |
| V | 10 | 1 | 130 |
| Total (Da) | | | 4589 |

The molecular weight of [^13^C,^15^N]-Sup35NM is 30084.9 Da so the carbonyl weighing factor is:

4589 Da of ^13^C carbonyl carbons / 30084.9 Da in the protein = 15.2%

To determine how many mg Sup35NM are present at various concentrations, we use the extinction coefficient of [^13^C,^15^N]-Sup35NM, ε = 0.8509 (mg/mL)^-1^ * cm-^1^ and the sample volume. For example:

25 µM * 30 µL * 0.8509 (mg/mL)^-1^ * cm-^1^ = 0.03 mg.

To determine how many µg of carbonyl carbons are present in the [^13^C,^15^N]-Sup35NM, we multiply by the carbonyl weighing factor:

30 µg * 15.2% = 4.6 µg of carbonyl carbons in Sup35NM at 25 µM.

To determine the specificity for the protein of interest, we divided of the amount of labeled carbonyl carbons in the protein of interest by the total number of labeled carbonyl carbons in the sample.

4.6 µg (in Sup35NM at 25 µM) / (4.6 µg in Sup35NM)+(5.7 µg in yeast lysates) = 45%
